# Transcriptional repression by ApiAP2 factors is central to chronic toxoplasmosis

**DOI:** 10.1101/100628

**Authors:** Joshua B. Radke, Danielle Worth, Dong-Pyo Hong, Sherri Huang, William J. Sullivan, Emma H. Wilson, Michael W. White

## Abstract

Bradyzoite differentiation is marked by major changes in gene expression resulting in a parasite that expresses a new repertoire of surface antigens hidden inside a modified parasitophorous vacuole called the tissue cyst. The factors that control this important life cycle transition are not well understood. Here we describe an important *Toxoplasma* transcriptional repressor mechanism controlling bradyzoite differentiation that operates exclusively in the tachyzoite stage. The ApiAP2 factor, AP2IV-4, is a nuclear factor dynamically expressed in late S phase through mitosis/cytokinesis of the tachyzoite cell cycle. Remarkably, deletion of the AP2IV-4 locus resulted in the increased expression of bradyzoite mRNAs in replicating tachyzoites, and in two different genetic lineages we confirmed the misexpression of tissue cyst wall components (e.g. BPK1, MCP4, CST1) and the bradyzoite surface antigen SRS9 in the tachyzoite stage. In the murine animal model, the loss of AP2IV-4 had profound biological consequences. Type II prugniaud strain parasites lacking AP2IV-4 were unable to form tissue cysts in brain tissue and the absence of this factor also recruited a potent immune response characterized by increases inflammatory monocytes, IFN-γ and higher numbers of both CD8+ and CD4+ T-cells. Altogether, these results indicate that suppression of bradyzoite antigens by AP2IV-4 during acute infection is required for *Toxoplasma* to establish a chronic infection in the immune-competent host.

**Author Summary:** The *Toxoplasma* biology that underlies the establishment of a chronic infection is developmental conversion of the acute tachyzoite stage into the latent bradyzoite-tissue cyst stage. Despite the important clinical consequences of this developmental pathway, the molecular basis of the switch mechanisms that control formation of the tissue cyst is still poorly understood. A fundamental feature of tissue cyst formation is the expression of bradyzoite-specific genes. Here we show the transcription factor AP2IV-4 directly silences bradyzoite mRNA and protein expression in the acute tachyzoite stage demonstrating that developmental control of tissue cyst formation is as much about when not to express bradyzoite genes as it is about when to activate them. Loosing the suppression of bradyzoite gene expression in the acute tachyzoite stage caused by deleting AP2IV-4 blocked the establishment of chronic disease in healthy animals through the pre-arming of the immune system suggesting a possible strategy for preventing chronic *Toxoplasma* infections.

## Introduction

*Toxoplasma gondii* is an obligate intracellular parasite that exhibits a multi-host and multi-stage developmental life cycle. Sexual stages are restricted to the gut mucosa of the feline definitive host and asexual stages of the intermediate life cycle occur within any warm-blooded host, including humans. Acute disease is generally asymptomatic in immune-competent hosts, however, primary or recrudescent infection from latent bradyzoites in humans with AIDS, those undergoing chemotherapy or in the unborn cause significant disease and death [1]. While the tachyzoite lytic cycle is responsible for disease pathology in human hosts, the interconversion of the tachyzoite stage into the bradyzoite stage underlies chronic infection and ensures host to host transmission [2]. Evidence indicates that unknown mechanisms regulating the tachyzoite cell cycle are intricately tied to bradyzoite differentiation with the choice to continue tachyzoite replication or develop into the latent bradyzoite containing tissue cyst made during S phase and mitosis of the immediate prior cell cycle [3, 4].

Transcriptome data demonstrates that unique changes in mRNA expression occur in the tachyzoite cell cycle and during development [5–7]. An estimated ~5% of all transcripts are exclusive to a single developmental stage with nearly 40% of the mRNAs in the tachyzoite division cycle periodically expressed. How these changes are controlled is largely unknown. Early genome mining for known gene specific transcription factors revealed two important observations. While the general transcriptional machinery is present in the genomes of Apicomplexa species, initial studies failed to identify classic gene specific transcriptional regulators common to higher eukaryotes. Second, an overall lack of genes encoding DNA binding proteins suggested a limited arsenal from which to regulate these dynamic changes in parasite developmental gene expression. In 2005, a family of DNA binding proteins (ApiAP2 factors) distantly related to the APETALA family plant transcription factors was discovered encoded in the genomes of Apicomplexa species providing the first evidence for gene specific transcription factors to date [8]. In *Plasmodium* spp., ApiAP2 factors bind DNA with distinct sequence specificity [9, 10] via a novel domain swapping mechanism [11] and have non-transcriptional roles in sub-telomeric chromosome biology [12]. Examples of ApiAP2 gene-specific functions in *Plasmodium falciparum* are ookinete (AP2-O) and sporozoite (AP2-Sp) ApiAP2s that serve as stage specific transcriptional activators regulating motile stages within the definitive host mosquito [13, 14]. Genetic disruption of AP2-O results in non-invasive ookinetes [14] whereas disruption of the AP2-Sp locus yields a parasite unable to form viable sporozoites [13]. The *Toxoplasma* genome encodes 67 ApiAP2 domain-containing proteins (ToxoDB and ref. [15]), with 24 of these genes expressed cyclically during the tachyzoite division cycle [5]. Interestingly, the peak mRNA expression of the cell cycle regulated ApiAP2s forms a time order sequence that spans the complete division cycle providing a mechanism to provide “just in time” delivery of gene products throughout the cycle [5, 16]. In *Toxoplasma*, ApiAP2s have been implicated in virulence and invasion mechanisms [17], as part of chromatin remodeling complexes [18, 19] and RNA processing machinery [20] and there is evidence for ApiAP2 factors regulating bradyzoite development. AP2XI-4 is up-regulated during bradyzoite development and the loss of AP2XI-4 blocks the stress-induction of some bradyzoite mRNAs, including the canonical marker, BAG1 [21]. The novel stress-inducible transcriptional repressor AP2IX-9 acts to prevent premature commitment to bradyzoite development through direct interaction with bradyzoite specific promoters [22].

Here we describe the discovery of a new level of developmental control in the intermediate life cycle that is required to establish the chronic tissue cyst stage in animals. AP2IV-4 is exclusively expressed in the tachyzoite division cycle with peak expression of the encoded mRNA and protein during early mitosis. Surprisingly, genetic knockouts of AP2IV-4 demonstrate it is non-essential to the replicating tachyzoite but is instead critical for the suppression of bradyzoite surface antigens and cyst wall proteins in the tachyzoite stage. Results from animal studies determined that AP2IV-4 silencing of bradyzoite gene expression is critical to enable tachyzoites to escape an effective immune response and produce the tissue cysts required for transmission.

## Results

### Many ApiAP2 factors dynamically regulated in the tachyzoite cell cycle are nonessential for growth

Previous studies identified ApiAP2 genes that are cyclically transcribed once per tachyzoite cell cycle with the peak timing of mRNA levels distributed throughout the division cycle [5]. The functions of periodically expressed ApiAP2 factors is largely unknown, although it is proposed they control the remarkable “just-in-time” cell cycle transcriptome of asexual stage Apicomplexa parasites [5, 16]. A Group-of-12 of these periodic ApiAP2 mRNAs share overlapping cyclical profiles that reach maximum expression during the S through mitotic phases (S/M) of the *Toxoplasma* tachyzoite cell cycle (Fig. 1A). We made multiple attempts to knockout each of the Group-of-12 ApiAP2 genes (Fig. 1B) in a Type I RH strain (RH*Δku80Δhxgprt*=RHQ strain) engineered for enhanced homologous recombination [23, 24]. The results from this series of genetic experiments were mixed; half the Group-of-12 ApiAP2 genes were successfully disrupted in the RHQ strain at relatively high prevalence except AP2III-2 (Fig. 1B), while knockouts of the other half failed repeated attempts. A recent whole genome CRISPR screen performed in human fibroblast cells (HFF)[25] supports the preliminary RHQ experimental sorting of Group-of-12 ApiAP2 genes into dispensable versus required (Fig. 1B, CRISPR column). Alternate developmental expression (and possible function), may help explain why half of the Group-of-12 ApiAP2s are not required for tachyzoite growth. AP2IX-4 (Huang et al, in preparation) and AP2XI-4 [21] are expressed in bradyzoites and recent studies indicate important roles for these factors in tissue cyst development; AP2III-2, AP2VI-1, and AP2XI-1 are also expressed in tachyzoites and bradyzoites [26]. Notably, AP2VI-1 mRNA is expressed at high levels across the *Toxoplasma* intermediate and definitive life cycles (the only ApiAP2 with this profile) and AP2III-2 is highly expressed in unsporulated oocysts [26]. The three Group-of 12 ApiAP2 factors exclusively expressed in tachyzoites (AP2IV-4, AP2XII-2, AP2XII-9)[26] also failed knockout attempts in RHQ and had significant negative phenotype scores in the HFF/CRISPR study (Fig. 1B).

**Figure 1:**
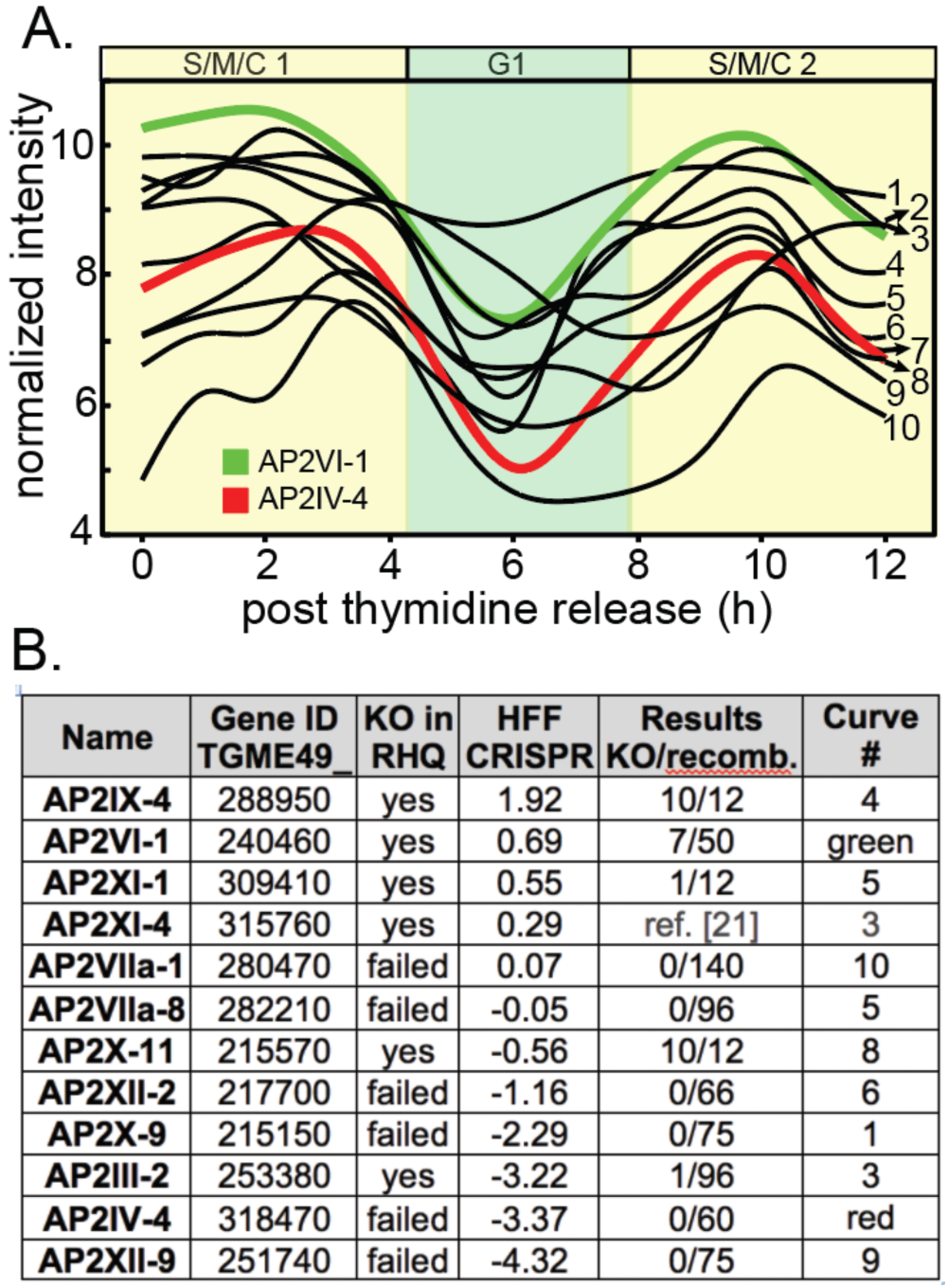
Genetic disruption of S/M regulated ApiAP2 genes. **(A.)** Shown are the cyclical mRNA profiles of twelve *Toxoplasma* ApiAP2s (Group-of-12) that reach maximum expression in the second half (S/M/C) of the tachyzoite cell cycle (results from our cell cycle transcriptome dataset [5]). Gene IDs for the curve numbers are indicated in (B.). AP2IV-4 is one of the most dynamic cyclical ApiAP2 mRNAs (~10 fold change, red curve), which peaks during late S phase to early mitosis and is down regulated by the start of G1 phase (see Fig. 2). The mRNA encoding AP2VI-1 (green curve) is similarly cell cycle regulated with peak expression slightly earlier than AP2IV-4 **(B.)** Multiple genetic knockout attempts in the RHQ strain were undertaken for eleven of the Group-of-12 ApiAP2 genes (AP2XI-4 knockout results see ref. [21]). Results: number of confirmed knockout clones (KO) versus total drug-resistant recombinants screened by custom PCR is indicated for each Group-of-12 gene. HFF CRISPR: average phenotype score for the Group-of-12 ApiAP2 genes in a recent whole genome CRISPR analysis [25] was included as a reference. In general, negative CRISPR phenotype scores indicate potential greater dependency for tachyzoite growth in HFF cells.

At the mRNA level AP2IV-4 stood out as one of the most dynamic of the Group-of-12 ApiAP2s with a greater than 10-fold change in mRNA abundance over a ~2 h period in the second half of the tachyzoite cell cycle (Fig. 1A, red curve). The failure to disrupt AP2IV-4 in RHQ parasites indicated an important function in tachyzoite replication. To verify AP2IV-4 protein is cell cycle regulated, we introduced three copies of the HA epitope tag in frame with the C-terminus of the AP2IV-4 coding region by genetic knock-in, which preserved the native promoter and genomic flanking contexts. The gene model for AP2IV-4 (http://toxodb.org/toxo/app/record/gene/TGME49_318470) predicts a single exon structure that encodes a large protein with a single AP2 DNA binding domain (Fig. S1A, diagram), which was verified by Western analysis of AP2IV-4^HA^ expressing parasites (Fig. S1A, gel right). As with previously tagged *Toxoplasma* ApiAP2s (*e.g.* ref. [5, 22]), AP2IV-4^HA^ in the RHQ strain localized exclusively to the nucleus (Fig. 2A) and was cell cycle regulated (Figs. 2A and S1B) with a timing similar to its mRNA expression profile (Fig. 1A, red curve). In an asynchronous tachyzoite population, AP2IV-4^HA^ was detectable in ~25% of vacuoles due to cell cycle periodicity. To pinpoint the exact cell cycle expression of AP2IV-4^HA^, co-staining with two daughter cytoskeleton markers was utilized to define the timing of initiation, accumulation and degradation of the AP2IV-4^HA^ fusion protein in comparison to the earlier expressing cyclical ApiAP2 factor (Fig. 1A, green curve), AP2VI-1^HA^, also produced by genetic knock-in. Antibodies for the inner membrane complex (Fig. S1B, α-TgIMC1)[27] and the apical cap (Fig. 2, α-TgISP1) [28] permit the late S phase through mitotic periods of the tachyzoite cell cycle to be resolved in time. AP2IV-4^HA^ first appeared in tachyzoites lacking internal daughter IMC structures as did AP2VI-1^HA^, although by first AP2IV-4^HA^ detection AP2VI-1^HA^ had already reached maximum expression in these parasites (Fig. S1B, a vs. b images). The detection of AP2IV-4^HA^ prior to internal daughter structures indicates initiation of expression in late S phase just prior to the start of mitosis and before nuclear division. The rapid accumulation of AP2IV-4^HA^ paralleled the formation of ISP1 rings of the daughter parasites (Fig. 2A, a-c images) and the growth of the IMC1 daughter scaffold (Fig. S1B, c,e images), while during these same cell cycle transitions AP2VI-1^HA^ declined rapidly to undetectable levels (Fig. 2B, d,e images; Fig. S1B, d,f images). AP2IV-4^HA^ was highly expressed throughout budding (distinct mitotic U-shaped nuclear morphology, DAPI staining, Fig. 2A) and began to disappear following nuclear division in late cytokinesis (Fig. 2A, images d,e) and was no longer detectable after resolution of the mother IMC or in any G1 parasite (data not shown).

**Figure 2:**
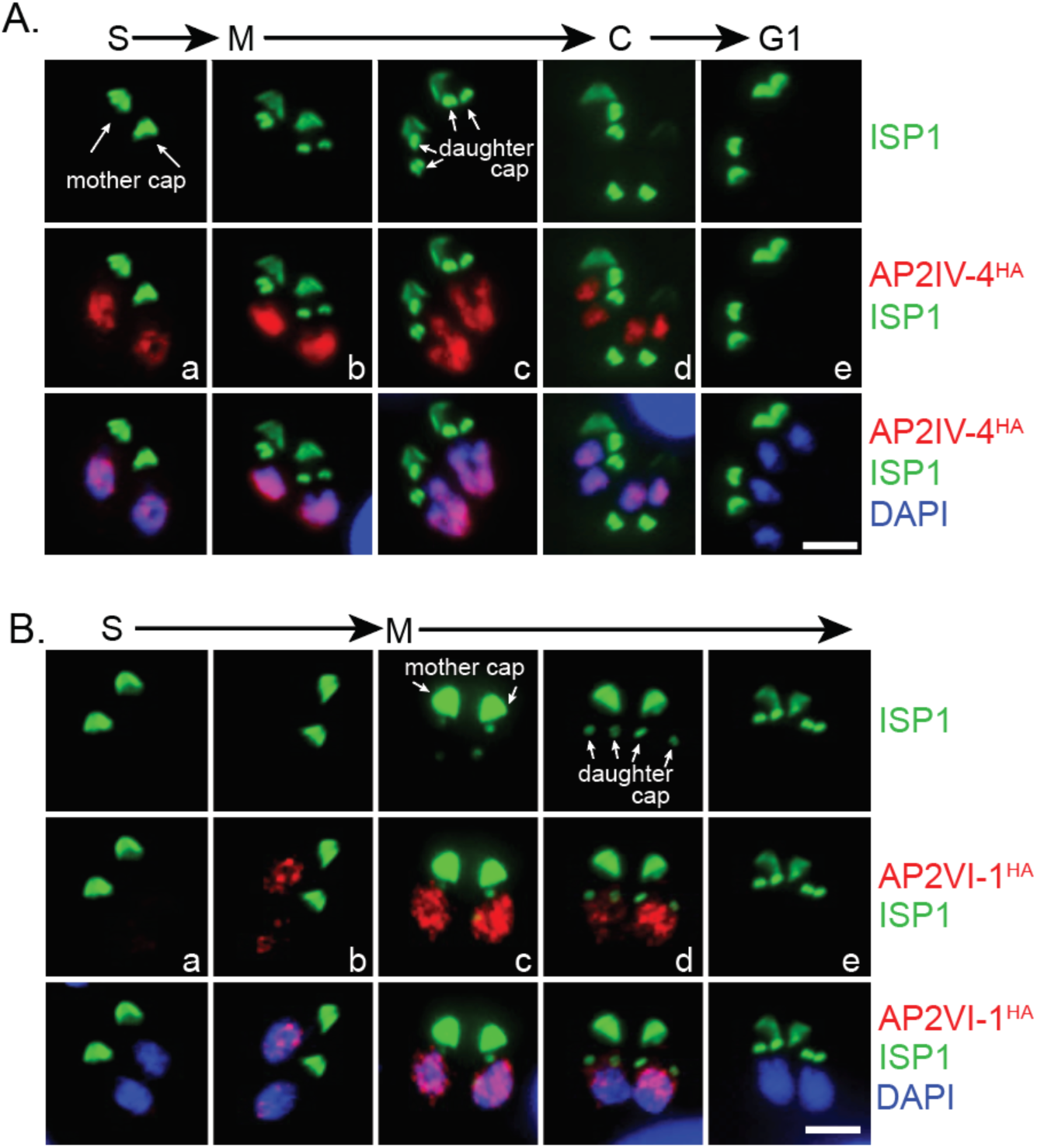
AP2IV-4 is dynamically regulated during the tachyzoite cell cycle. AP2IV-4 and AP2VI-1 were individually C-terminally epitope tagged with 3xHA at the endogenous loci of RHQ strain by genetic knock-in to preserve native promoter expression. Relative cell cycle progression for each experimental series is indicated about the images. **(A.)** AP2IV-4^HA^ expressing tachyzoites were co-stained with α-ISPI (apical cap, green) and DAPI (DNA, blue,) to determine the exact timing of AP2IV-4^HA^ expression (red) during the tachyzoite cell cycle. AP2IV-4^HA^ was first detected (image a) prior to daughter ISP1 formation and before the nucleus flattened in late S phase. Maximum AP2IV-4^HA^ protein (b, c) accumulation was marked by the appearance and elongation of daughter apical structures and U-shaped nuclear morphology. As the daughter apical cap (ISP1) matured and nuclear division was completed, AP2IV-4^HA^ disappeared near the end of cytokinesis (d). When all mother apical caps had been resolved and nuclear division completed (G1 phase), AP2IV-4^HA^ was not detectable. Scale bar=5 μm. **(B.)** AP2VI-1^HA^ parasites were also co-stained with α-ISP1 (apical cap, green) and DAPI (DNA, blue) to pinpoint AP2VI-1^HA^ expression (red) during the tachyzoite cell cycle. Initiation of AP2VI-1^HA^ expression also occurs prior to daughter apical cap formation (b) and achieves peak stabilization as daughter apical cap is formed (c). Apical cap rings (daughter caps, arrows) signify the degradation of AP2VI-1^HA^ expression (d) and AP2VI-1^HA^ is no longer detectable when the apical cap of the internal daughters is mature (e). Nuclear morphology restricts AP2VI-1^HA^ to S phase and early mitosis of the cell cycle, prior to the flattened or U-shaped nucleus of mitosis. Scale bar=5 μm

### Successful genetic knockout of AP2IV-4 indicates a nonessential role for tachyzoite growth

The failure to disrupt the AP2IV-4 gene in the RHQ strain (Fig. 1B) suggested this factor was essential for tachyzoite growth, although other explanations such as low frequency double crossover or growth defects preventing the recovery of AP2IV-4 knockout parasites could explain the knockout failure. To investigate whether low frequency recombination was responsible, we applied Cre-Lox methods [29] to disrupt AP2IV-4 using the rapamycin-inducible diCRE model recently introduced into the Type I RHΔ*hxgprt*Δ*ku80* strain (RHCre)[29]. To “flox” the AP2IV-4 gene in the RHCre strain, we performed serial epitope tagging by genetic knock-in of the AP2IV-4 (3xHA tag) and TGGT1_318480 (3xmyc tag) genes (see Fig. 3A diagram), which are in a sequential head to tail configuration on chromosome IV. The cell cycle expression of the AP2IV-4^HA^ fusion protein (data not shown) that resulted from the production of the RHCre-AP2IV-4^floxed^ strain was identical to the previously tagged AP2IV-4^HA^ protein in the RHQ strain (Fig. 2A). Gene TGGT1_318480 is expressed at very low levels in tachyzoites or bradyzoites (<30th mRNA percentile, ToxoDB.org) and was undetectable by both IFA and Western blot following tagging with 3xmyc. Cre-mediated excision of the AP2IV-4 locus was induced by a 6 h incubation with rapamycin (50nM) of RHCre-AP2IV-4^floxed^ parasites (Fig. 3A). In contrast to the failure to knockout AP2IV-4 in the RHQ strain by conventional methods (Fig. 1B), ~20% (10/51) of isolated clones following rapamycin treatment of RHCre-AP2IV-4^floxed^ parasites lacked the AP2IV-4 gene (also no longer HA+), and for two clones we verified the absence of AP2IV-4 mRNA (Fig. S2A). The isolation of viable RHCre-Δ*ap2IV-4* transgenic parasites indicated AP2IV-4 is not essential for tachyzoite growth.

**Figure 3:**
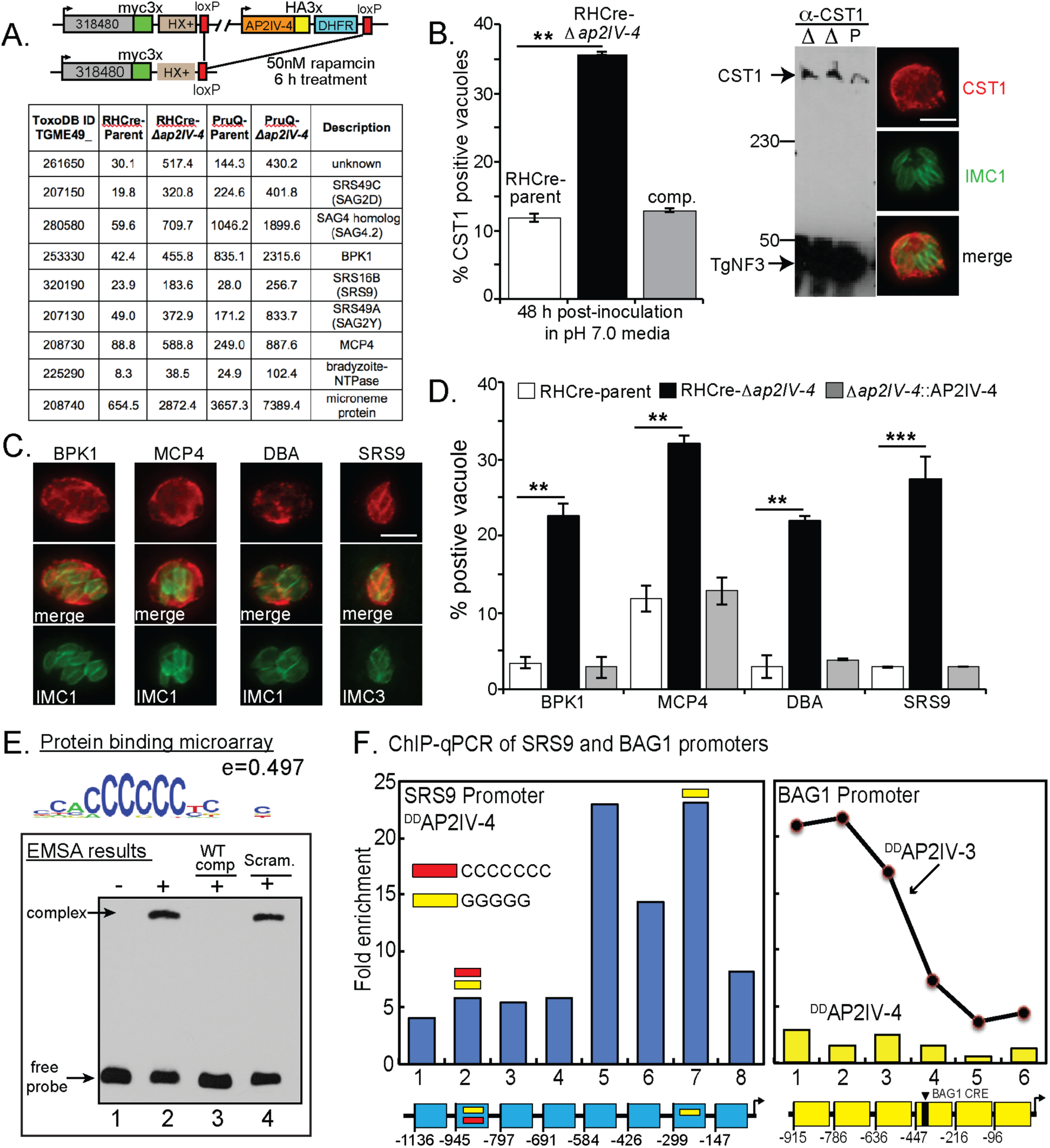
Tissue cyst wall and bradyzoite surface proteins accumulate in tachyzoites lacking AP2IV-4. **(A.)** Schematic representation of the AP2IV-4 knockout strategy in the RHCre strain. Insertion of loxP sites surrounding the AP2IV-4 locus was accomplished by sequential epitope tagging of TGGT1_318480 and AP2IV-4 genes utilizing the indicated tags and selectable markers. This resulted in the dual-tagged parental strain, RHCre-AP2IV-4^floxed^ (RHCre-parent). Addition of rapamycin as indicated activated the diCre-recombinase [29] thereby excising the AP2IV-4^HA^ tagged gene along with the pyrimethamine resistant DHFR gene. <u>Microarray analysis</u>: selected mRNAs altered by the knockout of the AP2IV-4 gene in RHCre-and PruQ-Δ*ap2IV-4* parasites; average RMA values for RHCre- or PruQ-parent (carrying AP2IV-4^floxed^) versus Δ*ap2IV-4* knockout transgenic strains are shown (complete results in Dataset S1). Note the higher baseline expression of mRNAs in the PruQ- versus the RHCre-parent strains. **(B.)** RHCre-parent (AP2IV-4^floxed^, open bar), -Δ*ap2IV-4* knockout (black bar), and-Δ*ap2IV-4*::AP2IV-4 complemented (grey bar) parasites were grown in HFF cells for 24 h under standard tachyzoite culture conditions (pH 7.0) and then IFA analysis performed using α-IMC1 (green stain, tachyzoite replication marker) and antibodies to cyst wall protein, CST1 (red stain). Scale bar=5 μm. CST1 positive vacuoles in each strain were quantified in triplicate by counting 100 vacuoles in randomly selected microscopic fields (**, p<0.01). Increased expression of CST1 protein in two independent RHCre-Δ*ap2IV-4* clones (Δ clones, see also Fig. S1A) compared to the RHCre-parental (P=AP2IV-4^floxed^) strain revealed by Western analysis. Nucleolar TgNF3 (43kDa) protein was included as a loading control [64]. Protein mass markers 230kDa and 50 kDa on left. Immunofluorescence images included on right are representative of CST1 positive vacuoles showing cyst wall localization in RHCre-Δ*ap2IV-4* parasites. **(C.)** Representative IFA images of RHCre-Δ*ap2IV-4* tachyzoites misexpressing bradyzoite proteins BPK1 and MCP4 with localization at the periphery of the vacuole consistent with DBA+ cyst wall structures, whereas, misexpressed bradyzoite-specific SRS9 was localized to the parasite surface. Note that SRS9 expression was always uniform for tachyzoites sharing the same vacuole as shown. Scale bar=5 μm. **(D.)** Increased numbers of BPK1, MPC4, DBA, and SRS9 positive vacuoles in RHCre-Δ*ap2IV-4* tachyzoites compared to RHCre-parent. Statistical significance indicated (**, p<0.01; ***, p<0.001). IFA analysis of RHCre-Δ*ap2IV-4*:AP2IV-4 (comp.) complemented parasites also shown. Positive staining vacuoles as in (C.) for each strain indicated was quantified in duplicate by counting 100 vacuoles in randomly selected microscopic fields. Note: IFA analysis and quantification of these bradyzoite proteins in PruQ-Δ*ap2IV-4* parasites is presented in Supplemental Materials, Figure S3. **(E.)** To determine whether the single AP2 domain in AP2IV-4 is capable of binding DNA, a protein binding microarray screen was completed using recombinant GST-AP2IV-4 protein (AP2 domain only)(see Material and Methods). The consensus binding sequence from the analysis was determined to be homopolymeric poly(dC):poly(dG); (5’-ACCCCCCT-3’/3’-TGGGGGGA-5’; enrichment score=0.497) the representative sequence logo shown indicates position weight matrices compiled for each base. An electrophoretic mobility shift assay was completed using 50ng of GST-AP2IV-4 protein and 20 fmol of a 59bp biotinylated DNA probe containing one 6 and one 5 homopolymer poly(dC) stretch (36% total poly(dC) nucleotides). A slowly migrating complex was identified (lane 2, top arrow) that was specifically competed by excess unlabeled poly(dC) competitor DNA (lane 3) but not by excess scrambled DNA fragment containing no more than three consecutive poly(dC) nucleotides (18% poly(dC) nucleotides). **(F.)**^DD^AP2IV-4 occupies the native SRS9 promoter in parasite chromatin (see Material and Methods for ChIP-qPCR assay details). Specific ^DD^AP2IV-4 binding was determined in eight specific regions of the SRS9 promoter 5’ to the SRS9 coding region (-1,136 bp flanking including the SRS9 5’-UTR up to the coding ATG). The region of the SRS9 promoter between -584 bp and -192 bp (regions 5-7) showed the highest specific enrichment for ^DD^AP2IV-4 binding. The occurrence of poly(dC) or poly(dG) motifs (5’-3’) in the SRS9 promoter are indicated by red or yellow bars, respectively. ChIP-qPCR analysis of ^DD^AP2IV-4 binding to the bradyzoite-specific BAG1 promoter in tachyzoite chromatin was included here as a negative control. A previous study established that the stress-induced ApiAP2 activator, AP2IV-3, specifically binds the BAG1 promoter [26] and is shown here only as a positive control reference. Note that the functionally mapped BAG1 CRE [65] is indicated in the included diagram (yellow diagram). All SRS9 and BAG1 primer sets used in the ChIP-qPCR studies are listed in Dataset S3.

### Microarray analysis reveals AP2IV-4 controls bradyzoite mRNA expression

Absent a requirement for AP2IV-4 in cell division, it was not clear what role AP2IV-4 serves in the tachyzoite. In order to build clues to understand AP2IV-4 function, duplicate total RNA samples from the RHCre-AP2IV-4^floxed^ parent and RHCre-Δ*ap2IV-4* tachyzoites were isolated (standard tachyzoite conditions, pH 7.0), converted to cRNA and hybridized to a custom *Toxoplasma* Affymetrix GeneChip [30]. In total, 40 mRNAs were altered >2-fold in RHCre-*Δap2IV-4* tachyzoites, including 26 mRNAs that were up-regulated (Fig. 3A, results of selected genes; complete results in Dataset S1). Remarkably, the loss of AP2IV-4 caused increased expression of mRNAs encoding known bradyzoite surface antigens (*e.g.* SRS9, SAG4.2) [31, 32] and cyst wall components (*e.g.* BPK1, MCP4)[6, 33] in the tachyzoite. These results indicated the major function for AP2IV-4 is to repress the transcription of key bradyzoite genes in replicating tachyzoites. This new level of developmental control of bradyzoite gene expression in the tachyzoite is distinct from the stress-induced AP2IX-9 mechanism we described previously [22]. Importantly, AP2IV-4 and AP2IX-9 combined appear to transcriptionally silence 73% (16/22, Dataset S2) of the bradyzoite genes thought to be activated by AP2XI-4 [21], which is also one of Group-of-12 ApiAP2s. The RHCre-Δ*ap2IV-4* transgenic strain was complemented with the cosmid PSBM794 [34] that carries a *Toxoplasma* genomic DNA fragment spanning the AP2IV-4 gene and RNA samples from the resulting RHCre-Δ*ap2IV-4*::AP2IV-4 transgenic strain were analyzed on the *Toxoplasma* GeneChip. The results from this experiment determined that reintroduction of the AP2IV-4 gene restored mRNA repression to RHCre-AP2IV-4^floxed^ parent levels for >80% of the mRNAs with the remaining mRNAs substantially reduced from the de-repressed levels of RHCre-Δ*ap2IV-4* tachyzoites (Dataset S1).

To confirm the function of AP2IV-4 in a second genetic lineage, we employed the same double-tagging strategy to "flox" the AP2IV-4 gene in the Type II Prugniaud strain (PruQ=Pru-Δ*ku80*Δ*hxgprt*)[35] followed by knockout of the AP2IV-4 gene by transient transfection of GFP-Cre (Fig. S2B, diagram)[36]. Two confirmed PruQ-Δ*ap2IV-4* clones lacking the AP2IV-4 gene were recovered from 178 independent clones screened (Fig. S2C). The successful disruption of the AP2IV-4 locus in PruQ confirms the dispensability of this factor for tachyzoite growth in a second genetic lineage. Microarray analysis of PruQ-Δ*ap2IV-4* tachyzoites identified very similar gene expression changes to RHCre-Δ*ap2IV-4* parasites; >90% of genes altered up or down by the loss of AP2IV-4 in these knockout strains were shared (Fig. 3A; complete lists Dataset S1). In the PruQ-Δ*ap2IV-4* parasites, bradyzoite mRNA fold changes were often less than in the RHCre-Δ*ap2IV-4* tachyzoites due to higher starting baseline levels of mRNA expression in the PruQ-AP2IV-4^floxed^ parent strain (Fig. 3A and Dataset S1). Type II Pru strains have significant capacity to spontaneously form bradyzoites (10−20%) in tachyzoite cultures (Fig. S3A and ref. [26]) and these spontaneously formed bradyzoites raise the population baseline expression of bradyzoite mRNAs. Nonetheless, microarray studies of PruQ-Δ*ap2IV-4* parasite mRNA expression clearly validated the conclusion that a major function of AP2IV-4 is to silence bradyzoite surface and cyst wall gene transcription in replicating tachyzoites.

### Tachyzoites lacking AP2IV-4 misexpress cyst wall and bradyzoite surface proteins

To verify bradyzoite proteins are misexpressed in replicating tachyzoites lacking AP2IV-4, we completed immunofluorescence assays (IFA) of RHQ-Δ*ap2IV-4* tachyzoites (Fig. 3B-D) followed by PruQ-Δ*ap2IV-4* tachyzoites (Fig. S3A,B) using antibodies to four bradyzoite specific proteins (α-BPK1, α-MCP4, α-CST1, α-SRS9) and also evaluated the formation of cyst walls using biotin-labeled *Dolichos biflorus* agglutinin (DBA). Microarray probes for the recently discovered bradyzoite cyst wall protein CST1 [37] were not included in *Toxoplasma* GeneChip, although we suspected the CST1 gene could be a target of AP2IV-4 suppression in tachyzoites. This was confirmed at the mRNA level by RT-qPCR using CST1-specific primers (not shown, see Dataset S3 for CST1 primer designs). Staining of RHCre-Δ*ap2IV-4* and PruQ-Δ*ap2IV-4* tachyzoites with CST1 antibodies confirmed increased expression (Figs. 3B and S3A) of this large cyst wall protein (>250 kDa), which was reversed by complementation of the RHCre-*Δap2IV-4* strain (Fig. 3B). Similar to CST1, the cyst wall pseudokinase BPK1 and structural protein MCP4 as well as bradyzoite surface protein SRS9 were all increased in RHCre-and PruQ-Δ*ap2IV-4* tachyzoites (Fig. 3C,D and Fig. S3A,B) as was the number of DBA+-vacuoles. Type I RH parasites are known to be resistant to developmental induction [38], and therefore, it was remarkable that deletion of single ApiAP2 factor could accomplish what strong alkaline stress poorly does in this strain. Further, misexpression of bradyzoite mRNAs in Δ*ap2IV-4* tachyzoites was similar to the levels of alkaline-induced expression (data not shown) and the encoded proteins were properly directed to either cyst walls or parasite surface (Figs. 3C and S3B) demonstrating that the loss of AP2IV-4 disrupted the timing not the localization or normal range of abundance. The misexpression of the bradyzoite antigens in Δ*ap2IV-4* populations was not 100% for either strain, which likely reflects the restricted S/M cell cycle window that AP2IV-4 operates (Figs. 1 and 2). The S/M timing of increased baseline bradyzoite mRNA expression in synchronized tachyzoites [5] is a similar cell cycle pattern. This hypothesis was examined further by co-staining PruQ-Δ*ap2IV-4* and PruQ-AP2IV-4^floxed^ parent tachyzoites with α-SRS9 and α-centrin antibodies (Fig. S3D). This IFA analysis determined that the majority of SRS9+/PruQ-Δ*ap2IV-4* parasites possessed duplicated centrosomes (S/M phases) confirming that SRS9 misexpression was occurring primarily in the mitotic half of the tachyzoite cell cycle. Interestingly, in the 14.3% of PruQ-AP2IV-4^floxed^ parental parasites spontaneously expressing SRS9 here too most parasites possessed duplicated centrosomes (3% of parental parasites were G1 phase and SRS9+). Altogether, these results are consistent with the unique relationship between bradyzoite differentiation and tachyzoite mitosis that we discovered more than a decade ago [3].

### AP2IV-4 binds DNA with sequence specificity

ApiAP2 factors have been shown to regulate gene expression through the binding of target promoters in a sequence specific manner [9, 10, 12-14, 17, 22]. To assess DNA binding specificity for AP2IV-4, a GST-AP2IV-4 fusion protein (AP2 domain only) was expressed, purified and incubated on a microchip containing all possible 10-mer DNA fragments (Fig. 3E, protein binding microarray results). A resulting 8-nucleotide “consensus” sequence motif bound specifically by the GST-AP2IV-4 fusion protein contains homopolymeric poly(dC):poly(dG) (Fig. 3E, 5’-ACCCCCCT-3’/3’-TGGGGGGA-5’; enrichment score 0.497). Electrophoretic mobility shift assays (EMSA) using DNA probes containing poly(dC):poly(dG) repeats were used to validate the specificity of GST-AP2IV-4 binding (Fig. 3E, EMSA results)[22]. GST-AP2IV-4 bound biotin-labeled DNA probes that contained a single instance of the “consensus” PBM motif and a second five nucleotide poly(dC) segment (Fig. 3E, lane 2), and was successfully competed using 300x excess unlabeled poly(dC) competitor DNA (Fig. 3E, lane 3), but not with unlabeled DNA probes that contained no poly(dC) stretch greater than three nucleotides (Fig. 3E, lane 4). In a larger protein binding screen of 46 *Toxoplasma* AP2 domains, two other ApiAP2 factors, AP2VIIa-5 and AP2XII-4, were determined to also specifically bind homopolymeric poly(dC):poly(dG) DNA (Kim et al, in preparation). In addition, a recent analysis of nucleosome-free regions for enriched DNA motifs discovered poly(dC):poly(dG) repeats were preferentially found upstream of cell cycle and bradyzoite genes, such as SRS9 (Wang et al, in preparation).

The presence of poly(dC):poly(dG) repeats in the SRS9 promoter (Fig. 3F, blue legend) suggested AP2IV-4 might directly binding this promoter. To examine this question, we utilized the FKBP (DD)/Shield 1 conditional expression model [39] in order to improve the signal strength for AP2IV-4 expression, which has been very successful for studying ApiAP2 factors in *P. falciparum* and *Toxoplasma* [22, 40, 41]. The FKBP peptide combined with three copies of the HA epitope tag was fused to the N-terminus of the AP2IV-4 coding region (^DD^AP2IV-4) by genetic knock-in methods. The addition of Shield 1 (100nM) to RHQ-^DD^AP2IV-4 transgenic parasites successfully increased nuclear levels of ^DD^AP2IV-4, but did not disrupt the normal periodic cell cycle expression of this protein (data not shown). Thus, there are likely significant post-transcriptional mechanisms regulating AP2IV-4 expression in tachyzoites as we also documented for AP2IX-9 expression previously [22]. Utilizing lysates prepared from RHQ-^DD^AP2IV-4 tachyzoites incubated with Shield 1, we performed chromatin immunoprecipitation followed by quantitative PCR of eight regions covering ~1,200bp of the SRS9 promoter and 5’-UTR (Fig. 3F). The results from this experiment showed that binding of ^DD^AP2IV-4 to the SRS9 promoter in parasite chromatin was enriched in regions 5-7 that includes a poly(dG) motif (region 7, yellow bar) ~230bp upstream of the SRS9 ATG (Fig. 3F). To control for non-specific binding, we analyzed ^DD^AP2IV-4 binding to the chromosome region (-950 bp) 5’-flanking of the BAG1 bradyzoite gene. In contrast to stress-induced AP2IV-3, which we have recently reported activates BAG1 [26], AP2IV-4 does not regulate BAG1 (Dataset S1). No enrichment of ^DD^AP2IV-4 binding was detected to the six regions tested within the BAG1 promoter (Fig. 3F), whereas ^DD^AP2IV-3 binding to this promoter is significantly enriched [26] (Fig. 3F, reference line graph).

### The loss of AP2IV-4 prevents tissue cyst formation in vivo

PruQ-Δ*ap2IV-4* parasites have typical tachyzoite characteristics; unlike bradyzoites they grew synchronously within a shared vacuole and retained normal SAG1 surface antigen expression even as they misexpressed SRS9 (Fig. S3C). Shifting PruQ-Δ*ap2IV-4* parasites into alkaline media (pH8.2) effectively slowed growth and induced high levels of DBA+ tissue cysts (Fig. 4B, In vitro=93.1%). By contrast, infections of BALB/c mice with PruQ-Δ*ap2IV-4* parasites failed to produce tissue cysts in brain tissue (Fig. 4A). In addition, inoculation of PruQ-Δ*ap2IV-4* parasites into BALB/c mice resulted in less virulence compared to the PruQ-AP2IV-4^floxed^ parent strain (LD100 >10^6^) such that all mice were able to survive an inoculation of 10 million PruQ-Δ*ap2IV-4* parasites (Fig. 4B). All mice were confirmed to be infected by serology demonstrating there are distinct phenotypes for PruQ-Δ*ap2IV-4* parasites *in vitro* versus *in vivo*. The lack of tissue cyst formation and decreased virulence *in vivo* was correlated with a lower parasite burden measured in BALB/c mice at 6 days post-inoculation (Fig. 4C), although lower parasite burden was not due to an inability of PruQ-Δ*ap2IV-4* parasites to replicate in mice as evident in 5 day peritoneal exudates (Fig. 4D). As there was no evidence of attenuated parasite growth *in vitro* or *in vivo* (Fig. 4D), these results pointed to increased parasite control by the host.

**Figure 4:**
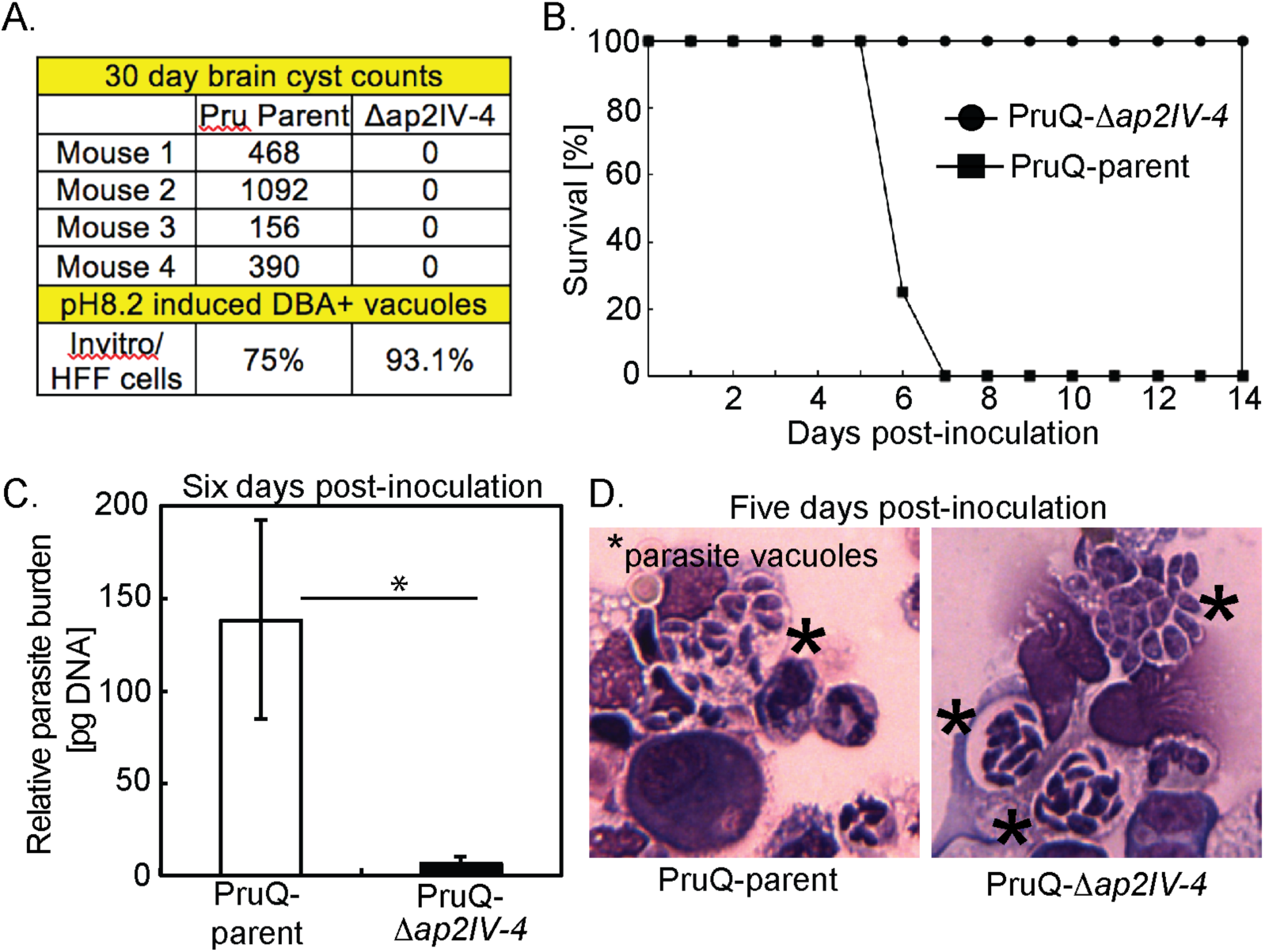
Loss of AP2IV-4 reduces tissue cyst number and increases animal survival. **(A.)** PruQ-*Δap2IV-4* parasites were induced by alkaline-stress (72 h, pH 8.2 media) to form tissue cysts (DBA+) at high levels in HFF cells, whereas, they were unable to form tissue cysts in brain tissue of BALB/c mice compared to PruQ-AP2IV-4^floxed^ (defined here as PruQ-parent). **(B.)** Inoculation of BALB/c mice (10-12 weeks old) with PruQ-Δ*ap2IV-4* versus PruQ-parent parasites at high inoculum (1×10^7^) shows that disruption of the AP2IV-4 gene further attenuated the avirulent PruQ-parental strain. **(C.)** BALB/c mice were inoculated intraperitoneally with 1×10^7^ parasites of either the PruQ-parental or PruQ-Δ*ap2IV-4* strain. At day 6 post-infection peritoneal exudate cells were harvested, total DNA was extracted and parasite burden was determined following amplification of the B1 gene compared against a standard curve. Statistical significance (*, p<0.05) is indicated **(D.)** Animals were infected with the strains indicated and at day 5 post-infection peritoneal cells were harvested and spun onto cytospin slides. Slides were stained using HEMA3 staining kit and examined under a light microscope. Intracellular parasites are denoted by a star.

### Misexpression of bradyzoite proteins in tachyzoites recruits an effective host immune response

The development of bradyzoites and cysts *in vivo* is, at least partly, dependent upon immune factors. This is illustrated in immune deficient animals where bradyzoites fail to develop and mice succumb due to uncontrolled tachyzoite replication. The protective immune response is dominated by the recruitment of inflammatory monocytes and T cell production of IFN-γ [42]. Yet, the host immune response is unable to clear the bradyzoite and cysts persist for long periods in host tissues. The pay off between protection and the development of chronic infection is poorly understood. At day 6 post *Toxoplasma* infection neutrophils and inflammatory monocytes, distinguished by their expression of Ly6 surface antigens and distinct morphologies (Fig 5A,B), were present at the site of infection. Although neutrophils are important sources of IL-12, inflammatory monocytes are the key effector cell in controlling parasite replication [43]. Analysis of the proportion of these populations following infection of parasites with an intact or disrupted AP2IV-4 loci revealed striking differences. Consistent with the response to a high inoculum of parasites [43], infection with the PruQ-AP2IV-4^floxed^ parasites (PruQ-parent) induced the influx of neutrophils, outweighing inflammatory monocytes by a ratio of 5:1 (Fig. 5A,B, and D). By contrast, PruQ-Δ*ap2IV-4* infections led to a significant increase in the proportion of inflammatory monocytes (Fig. 5A). This was confirmed by cytospin, where large numbers of polymorphonuclear neutrophils can be seen in the peritoneal exudate wash of mice infected with PruQ-parental parasites and foamy monocytes observed with PruQ-Δ*ap2IV-4* parasite infection (Fig. 5B). Further, the absolute numbers of cells recruited to the site of infection was significantly greater in the absence of AP2IV-4 (Fig. 5B,C). By day 5 post infection, T cells have been activated and have migrated to sites of parasite replication [44, 45]. Consistent with an overall increase in cell infiltration following infection with PruQ-Δ*ap2IV-4* parasites, significantly more CD4 and CD8 T cells were observed in the peritoneum of mice infected with PruQ-Δ*ap2IV-4* parasites compared to infection with the PruQ-parental strain (Fig. 5E). Measurement of serum concentrations of IFN-γ over the course of infection revealed a significant increase at day 5 post infection that is resolved by day 6 (Fig. 5F). IFN-γ is the primary cytokine responsible for controlling parasite replication by activating infected cells. These data indicate that in the absence of AP2IV-4 silencing of bradyzoite gene expression in the tachyzoite stage there is an amplification of the protective immune response at the innate and adaptive levels.

**Figure 5:**
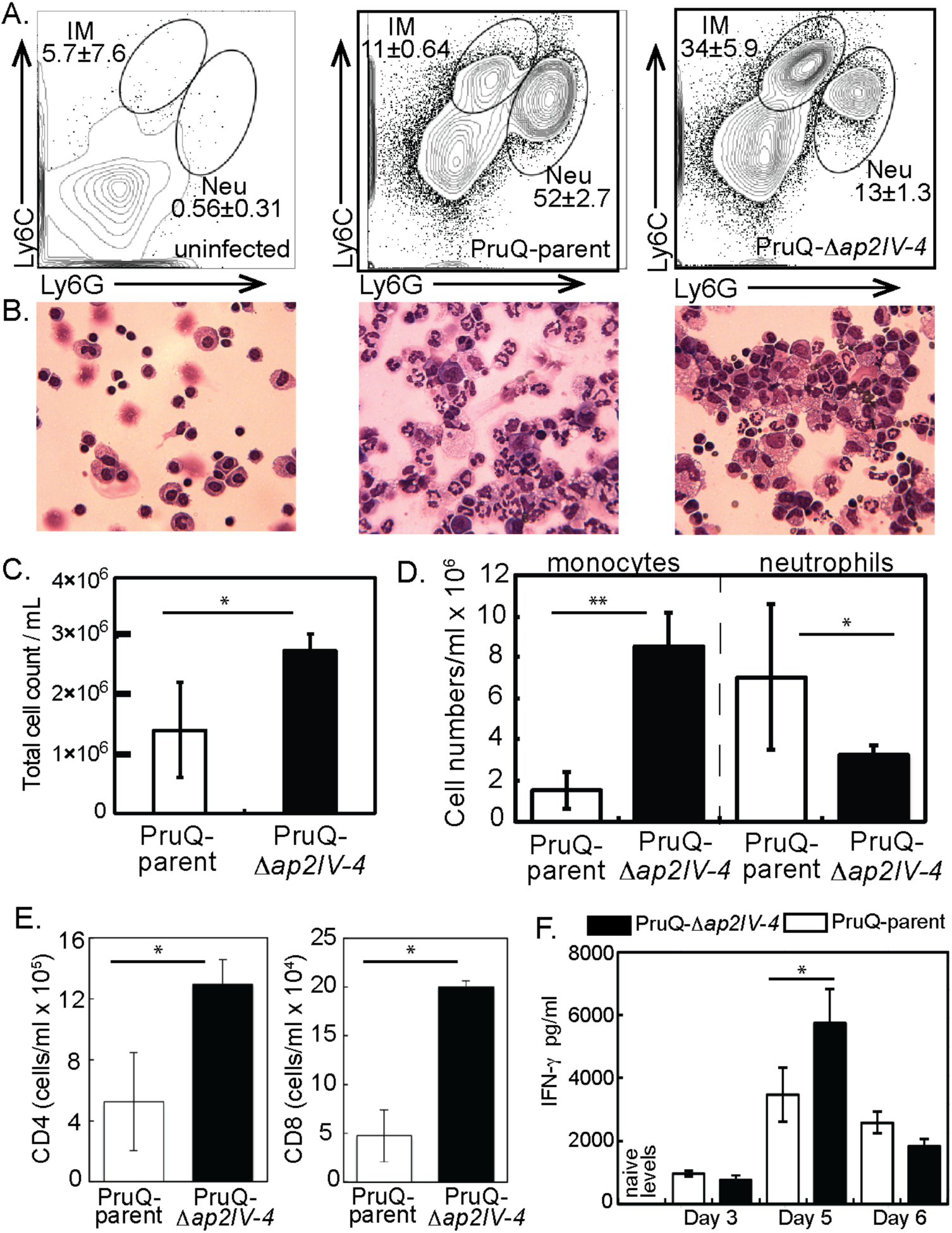
Infection with PruQ-*Δap2IV-4* tachyzoites results in greater cell infiltration to the peritoneum, and preferentially recruits inflammatory monocytes. **(A.)** Peritoneal exudate cells were harvested at day 6 post-infection from mice inoculated with PruQ-Δ*ap2IV-4* versus PruQ-AP2IV-4^floxed^ (PruQ-parent) parasites and subjected to immunostaining for flow cytometry. Cells were stained for extracellular antigens, and gated on CD45 positive cells for analysis. Cells were analyzed for expression of Ly6C and Ly6G. **(B.)** Representative cytospin images are provided immediately below the matching FACS plots above. **(C.)** Total cell infiltration into the peritoneum of infected mice. Cell counts were quantified based on volume recovered from the peritoneal wash. **(D.)** Absolute cell quantification of inflammatory monocytes and neutrophils. **(E.)** Quantification of CD3+CD4+ and CD3+CD8+ T cell populations recruited to the peritoneum. **(F.)** Serum was collected from infected animals at days 3, 5 and 6 post infection, and systemic levels of IFN-γ were quantified. Statistical significance (*, p<0.05; **, p<0.01) is indicated.

## Discussion

The life cycle of *Toxoplasma* is heteroxenous with a sexual definitive cycle in the felid host and a second intermediate life cycle in any endothermic animal including humans. The steps of the intermediate life cycle leading to tissues cysts in murine brain tissue illustrate this developmental process [4]. Bradyzoite/sporozoite oral infection leads to population wide development of the tachyzoite stage [46, 47] that is followed by systemic spread of tachyzoites. In particular, spread into the vasculature resulting in the infection of endothelial cells of brain capillaries is a critical route for tachyzoites to cross the BBB into the brain [48]. Through poorly understood mechanisms, the tachyzoites slow growth [4] and alter their transcriptome to form dormant bradyzoite-tissue cysts in neurons [49] setting the stage for transmission to the next host animal. Thus, there are two competing demands of the *Toxoplasma* intermediate life cycle; expand tachyzoite numbers to ensure systemic spread within a host [48] and produce the dormant bradyzoite-tissue cyst required for passing the infection onto a new host [4]. How *Toxoplasma* mechanistically balances these competing demands is not understood. However, clues to an answer are emerging from our studies of ApiAP2 factors (see Fig. 6 model). Early bradyzoite development is associated with the induction of six *Toxoplasma* ApiAP2 genes (AP2Ib-1, AP2IV-3, AP2VI-3, AP2VIIa-1, AP2VIII-4, AP2IX-9) that are not expressed in the tachyzoite [22, 26]. Remarkably, these factors do not operate in the same direction. AP2IX-9, is a stress-inducible repressor of bradyzoite gene expression [22], while AP2IV-3 is a stress-induced transcriptional activator (and likely AP2Ib-1) regulating many of the same bradyzoite genes as AP2IX-9 [26]. The studies here add unexpected new complexity to bradyzoite developmental gene expression. AP2IV-4 is the first tachyzoite-specific transcription factor we have identified that regulates the formation of tissue cysts. Thus, our studies have uncovered a complex ApiAP2 transcriptional network of repressors and activators competing at the interface of tachyzoite replication and early switching to regulate tissue cyst formation (see Fig. 6 model). Notably, we have not yet identified an ApiAP2 that exclusively operates late in bradyzoite development. For the ApiAP2 gene family, the most distinguishing feature of mature bradyzoites is the down regulation of many ApiAP2 factors [26].

**Figure 6:**
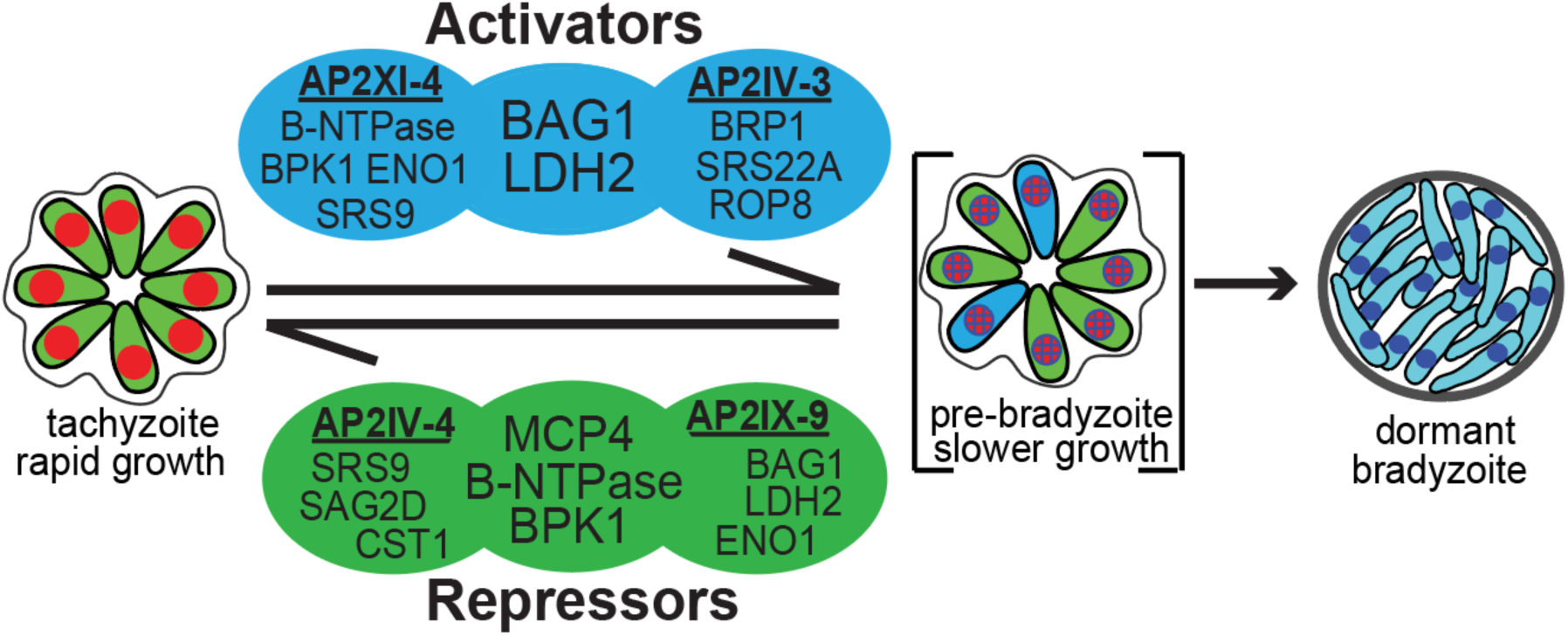
ApiAP2 transcriptional network controlling bradyzoite development. Transcriptional activators AP2XI-4 [21] and AP2IV-3 [26] co-regulate BAG1 and LDH2 genes and independently regulate other bradyzoite genes (representative genes shown, see Dataset S2 for complete gene lists). Similarly, transcriptional repressors AP2IV-4 and AP2IX-9 [22] regulate common (B-NTPase, MCP4, BPK1) and independent sets of bradyzoite genes. Note, AP2XI-4 and AP2IV-4 are Group-of-12 cell cycle ApiAP2s of tachyzoites (Fig. 1), while AP2IV-3 and AP2IX-9 are stress-inducible factors not expressed in replicating tachyzoites [22, 26].

Does the lack of ApiAP2 factors specific for mature bradyzoites mean that once initiated bradyzoite development *in vivo* progresses to maturity? Answering this question will be challenging given the asynchrony of bradyzoite development. However, a recent analysis of tissue cyst biology provides two important insights; tissue cyst size in the infected murine brain is related to tachyzoite vacuole size at the time of switching and average cyst numbers in tissues like murine brain become stable after an early period [50]. Add to this the observation that tissue cyst recrudesence is rare in the brain of immune-competent animals [51], returns the discussion to the critical importance of the tachyzoite stage for fulfilling the biotic demands of the intermediate life cycle. The discovery of tachyzoite-specific AP2IV-4 highlights the concept that specific life cycle decisions begin upstream in the developmental pathway and provides insight into the mechanisms that link the tachyzoite cell cycle to bradyzoite development. Together, the cell cycle AP2IV-4 and the stress-inducible AP2IX-9, comprise two independent levels of transcriptional regulation preventing bradyzoite development in *Toxoplasma*. This is convincing support for the hypothesis that tachyzoite growth is the primary driver of parasite biomass and through dissemination the tachyzoite finds suitable host cell environments for which to develop ultimately end-stage bradyzoites [4, 22]. In addition, the overlap of gene regulatory targets between AP2IV-4 and AP2IX-9 (Dataset S2) indicate there is some redundancy governing the induction of the bradyzoite developmental pathway indicating the importance of preventing premature commitment to the bradyzoite stage that leads to dormancy [22]. The earliest clues to unfolding bradyzoite development in *Toxoplasma* revolves around the central role asexual stage replication plays in the transition to growth-arrested end-stages [3, 52, 53]. DNA replication in the tachyzoite is required for bradyzoite development [52] and the tachyzoite is “poised” to enter the bradyzoite developmental pathway during each round of replication [3]. The sub-transcriptome of the tachyzoite S and mitotic phases is enriched in basal bradyzoite transcripts [5] and developing populations have more 2N parasites [3], which is a cell cycle timing that corresponds with peak AP2IV-4 expression. This places AP2IV-4 perfectly within the tachyzoite cell cycle to regulate these critical developmental processes. Repressing tissue cyst wall formation in the tachyzoite could provide the parasite with flexibility to maintain a replicative stage or quickly interpret “development” signals in the animal resulting in induction of bradyzoite differentiation when the parasite encounters the immune system and/or a tissue that favors tissue cyst longevity. In addition to controlling when and where tissue cyst formation occurs, repressors like AP2IV-4 may need to be re-expressed for pre-bradyzoites or bradyzoites to recrudesce. Consistent with this idea, our previous studies demonstrated bradyzoites from murine brain cysts re-express tachyzoite antigens prior to their first division in HFF cells and most bradyzoites that failed to re-express them did not divide [3].

These studies also demonstrate that deletion of a single ApiAP2 factor in *Toxoplasma* can significantly alter the course of the host immune response. Thus, host influences on ApiAP2 evolution has likely led to mechanisms that suppress bradyzoite antigens during acute infection, which we show here is required to establish a chronic infection in the murine brain. There are implications from this discovery for future vaccine development that might block tissue cyst formation in food animals, and thereby eliminate this source of human infections, which is an unmet therapeutic challenge. Our results point to three major components of the immune response that are increased in the absence of AP2IV-4; i) the recruitment of inflammatory monocytes [42, 54], ii) T cell recruitment, and iii) IFN-γ production (Fig. 5). A rapid response to the signature of a fast replicating lytic parasite is appropriate but there would be little evolutionary drive to respond equivalently to a slow replicating cyst form. Thus, changing the signatures of the parasite as we have done here with the deletion of AP2IV-4 have dramatically altered the early immune response, with bradyzoite antigens now being seen in the context of significant cell lysis. Increased inflammatory monocyte recruitment may point to a change in the ability of the parasite to be seen by the innate immune response. This could be either increased TLR recognition of bradyzoite antigens or a failure to inhibit by the tachyzoite. Perhaps predictably in the presence of enhanced monocyte recruitment, the T cell response is also superior. Alternatively, the misexpression of bradyzoite antigens in the replicating and systemic tachyzoites lacking AP2IV-4 may increase the overall function of the immune response by either targeting it more rapidly to the bradyzoite or act as an adjuvant to the overall anti-parasite response. Further studies will be needed to fully understand the molecular mechanism(s) responsible for the shift to a more protective immune response. It is worth noting that the functions now emerging for AP2IV-4 in controlling *in vivo* persistence were not uncovered by cell culture models. Achieving bradyzoite switching *in vitro* in the mid-90’s was a major breakthrough, and much has and will be learned using these models [55, 56].

However, the complexity of parasite encounters with host cells and tissues in animals can not be replicated by these models. Distinct tissue tropisms observed for tissue cyst formation in animals infected with *Toxoplasma* [46, 57–59] suggest the parasite senses different host cell environments and relays this information to the mechanisms controlling developmental switching [4]. We know little about the molecular basis for *Toxoplasma* host tissue tropisms, however, it is likely that the network of ApiAP2 repressors and activators we have discovered will have critical roles in these host-parasite interactions.

## Materials and Methods

A summary analysis of the AP2IV-4 microarray data can be found in Dataset S1. For a comparative analysis of genes regulated by AP2IV-3, AP2XI-4, AP2IV-4 and AP2IX-9, please see Dataset S2. All new gene expression data sets produced in this study have been uploaded to the Gene Expression Omnibus (GSE93531). All transgenic strains and oligonucleotides used in this study are found in Dataset S3

### Cell culture, genetic manipulation and conditional genome engineering

*Toxoplasma* tachyzoites were serially passaged *in vitro* using confluent tissue culture flasks (T25cm^2^ and T175cm^2^) containing human foreskin fibroblasts (HFF cells; obtained from ATCC, Manassas, VA). <u>Conventional ApiAP2 knockouts:</u> To create ApiAP2 gene replacement plasmids for Group-of-12 factors (Fig. 1), the 5’ UTR and 3’UTR regions (>500bp each flank) were PCR amplified from RH genomic DNA and cloned via the BP reaction into pDONRP4-P1R or pDONRP2R-P3, respectively. The pDONRP1-P2 plasmid containing the HXGPRT selectable marker is described [60]. Finally, the three entry plasmids (5’ApiAP2 UTR:HXGPRT:3’ApiAP2) were combined in the LR reaction with pDESTP4-P3 to create the expression plasmid pDEST"ApiAP2" HXGPRT KO. All BP and LR reactions were done according to manufacturers protocol (Thermo Fisher); 25μg of pDEST"ApiAP2" HXGPRT KO plasmid was transfected into 2.5×10^7^ RHQ tachyzoites then parasites were selected in media supplemented with xanthine (40mg/ml) and mycophenolic acid (50mg/ml). When drug resistant parasites were observed, clones were isolated by limiting dilution. <u>Introduction of loxP recognition sites:</u> loxP recognition sites were introduced by inverse PCR amplification of the pLIC-3xHA/DHFR and pLIC-3xmyc/HXGPRT plasmids, followed by Apal digestion of the linear PCR product and T4 DNA ligation to re-circularize the plasmid. This resulted in the introduction of a loxP site downstream of the indicated *T. gondii* selectable marker (ex. Fig 3A), creating the plasmids pLIC3xmyc/HXGPRT/loxP and pLIC-3xHA/DHFR/loxP. *Toxoplasma* gene TGGT1_318480 was fused with a triple repeat of the myc3 epitope in RHCre or PruQ by homologous recombination using the pLIC3xmyc/HXGPRT/loxP plasmid. Likewise, AP2IV-4 (TGGT1_318470) was fused with a triple HA epitope in clonal isolates of RHCre-318480^myc^ or PruQ-318480^myc^ by homologous recombination with the pLIC-3xHA/DHFR/loxP plasmid, thereby “floxing” the AP2IV-4 genomic locus. Rearrangement of the genetic locus of interest was verified by nested PCR for all clones isolated. Please see Dataset S3 for strain details and oligonucleotide sequences. <u>CRE-recombinase expression:</u> Expression of CRE-recombinase strain was induced in RHCre-AP2IV-4^floxed^ parasites as previously described [29]. To initiate CRE mediated excision of AP2IV-4, RHCre-AP2IV-4^floxed^ parasites were seeded at a 5:1 MOI in HFF monolayers, allowed to invade for 2 h, washed 3x with Hanks balanced salt solution (Corning) to remove all free floating parasites and followed by 6 h treatment with 50nM rapamycin. The monolayers were scraped, twice passaged through a 25ga needle, filtered to remove host debris and immediately cloned by limiting dilution to reduce competition from the background of "wild type" AP2IV-4 parasites. For PruQ-AP2IV-4^floxed^, 5×10^7^ parasites were transfected with 25μg of pmin-CRE-eGFP plasmid [36], allowed to recover for 24 h, scraped, force lysed by passage through a 25ga needle, filtered and immediately cloned by limiting dilution. <u>Cosmid complementation:</u> Cosmid isolates that span the AP2IV-4 locus were identified with the gBrowse function at www.ToxoDB.org. PSBM794 was selected and 10μg of cosmid DNA was transfected into 1x10^7^ RHCre-Δ*ap2IV-4* parasites and a stable population selected using two rounds of phleomycin (5mg/ml) selection. <u>Conditional expression of AP2IV-4:</u> The AP2IV-4 (TGGT1_318470) coding sequence was PCR amplified from RH genomic DNA with oligonucleotides that included in-frame MfeI/EcoRV sites, which were used to clone the PCR fragment into the pCTDDHA3x plasmid (Dr. Boris Striepen, University of Georgia). The resulting plasmid, pCTDDHA3x-AP2IV-4, contains an N-terminal fusion of the FKBP peptide (11.2 kDa) and a triple repeat of the HA epitope (4.4 kDa) fused in-frame with AP2IV-4 (^DD^AP2IV-4) that allows for ectopic conditional expression of the fusion protein using the small molecule Shield 1 [22]. The plasmid was transfected into the RHQ strain and transgenic parasites selected using chloramphenicol (34mg/mL) with individual clones isolated by limiting dilution. <u>Endogenous epitope fusions with 3xHA:</u> *Toxoplasma* genes AP2IV-4 (TGGT1_318470) and AP2VI-1 (TGGT1_240460) were tagged at the genomic locus with a triple copy of HA in RHQ by homologous recombination using the pLIC-3xHA/DHFR plasmid as previously described [24].

### Immunofluorescence assays and Western analysis

Parasites were grown in confluent HFF cells and prepared for immunofluorescence as previously described [61]. Primary antibodies were used at the following concentrations: HA (rat mAb 3F10, 1:500, Roche); ISP1 (mouse mAb clone 7E8, 1:2000, Dr. Peter Bradley, University of California, Los Angeles); IMC1 (mouse mAb, 1:1000, Dr. Gary Ward, University of Vermont); biotin-labeled *Dolichos biflorus* agglutinin (DBA) (Vector labs, CA,1:3000); BPK1, MCP4 and SAG1 (mouse polyclonal antibodies=pAbs, 1:1000, Dr. John Boothroyd, Stanford University,); CST1 (mouse pAb, Salmon E, 1:2000, Dr. Louis Weiss, Albert Einstein College of Medicine); SRS9 (rabbit pAb, 1:1000, Dr. Laura Knoll, University of Wisconsin). Secondary antibodies by Alexa or streptavidin conjugated secondary antibodies were used at a 1:1000 dilution. All images were collected with a Zeiss Axiovert microscope equipped with 100x objective. Statistical significance was calculated using the one-tailed t-test, p values as indicated.

Protein from 25×10^6^ parasites were isolated, purified and whole parasite lysates collected as previously described [61] and subjected to electrophoresis on a SDS-PAGE gel. After transfer to nitrocellulose, the blots were probed with primary antibodies for CST1 (mouse pAb, Salmon E, 1:2000, Dr. Louis Weiss, Albert Einstein College of Medicine) and the loading control TgNF3 (mouse pAb, 1:1000, Dr. Stan Tomavo, Pasteur Institute, Lille). Detection of the proteins was completed using HRP conjugated antibodies (Jackson ImmunoResearch) followed by chemiluminescence reaction for visualization.

### RNA microarray

Two independent biological replicates of total RNA were isolated from five RHCre transgenic strains (Dataset S3): RHCre-AP2IV-4^floxed^ clone ID6, RHCre-Δ*ap2IV-4* clones 27 and 30, and cosmid complemented populations (PSBM794; RHCre-Δ*ap2IV-4*::AP2IV-4) of each knockout. Likewise, two biological replicates were isolated from the following PruQ strain transgenics (3 total strains): PruQ-AP2IV-4^floxed^ clone C3, PruQ-Δ*ap2IV-4* clones 10 and 34. RNA quality for all strains was evaluated using the Agilent Bioanalyzer 2100 (Santa Clara, CA) and 500ng of total RNA was prepared for hybridization on the ToxoGeneChip as described [5]. The resulting data was analyzed using GeneSpring GX software (v11.5, Agilent) and all microarray data made available in the Gene Expression Omnibus (GSE93531).

### Protein expression and electrophoretic mobility shift assay (EMSA)

The AP2 domain (amino acids 782-854) of AP2IV-4 was cloned into pGEX4T3 and expressed as a GST-fusion protein. Following affinity purification on a glutathione column, purified GST-AP2IV-4 protein was subjected to protein binding microarrays as previously described [9, 22]. Complementary oligonucleotides were annealed to create 5’-biotinylated DNA probes of 59bp (WT) and 60bp (scrambled). All binding reactions contained 20fmol DNA probe and 50ng of GST-AP2IV-4 protein. Non-biotinylated “cold” competitor probe was added at 300x concentration. GST-AP2IV-4-DNA complexes were resolved on a 6% polyacrylamide PAGE gel, transferred to a nylon membrane and interactions visualized using the LightShift Chemiluminescent EMSA kit as described by the manufacturer (Thermo Fisher, Waltham, MA). Oligonucleotide sequences used for GST-AP2IV-4 expression and DNA probes can be found in Dataset S3.

### Chromatin Immunoprecipitation and qPCR

Chromatin immunoprecipitation followed by quantitative PCR (ChIP-qPCR) was performed as previously published (supplement of ref. [22]). In brief, RH-^DD^AP2IV-4 and RHΔ*hxgprt* (negative control) parasites were inoculated at 3:1 MOI in T175 cm^2^ flasks, allowed to invade for 1 h, rinsed three times with Hanks balanced salt solution (Gibco) to remove free floating parasites and fresh media containing 100nM Shield 1 was added. Parasite cultures were allowed to grow 36 h prior to intracellular crosslinking with formaldehyde and isolation of nuclear fraction. Nuclear material was subjected to sonication to shear DNA into 200-1000bp fragments and soluble fraction incubated with α-HA antibody (5μg, ab9110, rabbit, Abcam). Protein-DNA complexes were isolated using protein-G coupled magnetic beads (Dynabeads, Invitrogen) and DNA isolated by treatment with 1% SDS followed by phenol-chloroform extraction and ethanol precipitation. Whole genome amplification (Sigma-Aldrich) was performed on ChIP-DNA and purified by Qiagen Mini-Elute PCR kit. qPCR was performed using 20ng/rxn of specific (^DD^AP2IV-4) chromatin and non-specific chromatin (RHΔ*hxgprt*) using Fast SYBR green master mix on an ABI7900 according to manufacturers protocols. Relative enrichment was calculated with the equation: 2^^-(_Δ_Ct target-_Δ_Ct non-target)^ where the change in Ct value of specific versus nonspecific chromatin at the SRS9 and BAG1 promoters was calculated. All ChIP-qPCR oligonucleotides used can be found in Dataset S3.

### Animals and Infections

Mice were purchased from Jackson or Harlan Laboratories. 10-12 week old BALB/c mice were infected with 1×10^7^ PruQ-AP2IV-4^floxed^ parental strain or PruQ-Δ*ap2IV-4* intraperitoneally in sterile PBS. Uninfected, age-matched mice were used as naïve uninfected controls. Mice were monitored daily and euthanized at day 6-post infection for study.

To examine acute virulence, 5-6 week old female BALB/c mice were injected intraperitoneally with 10^5^, 10^6^, or 10^7^ PruQ-AP2IV-4^floxed^ or PruQΔ*ap2IV-4* parasites (4 mice per group, 10^7^ dose only shown in Fig. 4). Plaque assays were performed for each sample and ensured equal viability between strains. Mice were examined daily and time to death was recorded. Serology performed on cardiac bleeds of infected mice confirmed presence of *Toxoplasma*. To assess cyst burden, mice were infected with 10^6^ parasites as described above and allowed to progress to chronic infection for 30 days (4 mice per group). Brains were then homogenized; homogenates were fixed, quenched, and permeabilized. Samples were blocked in 3% BSA/1xPBS/0.2% Triton X-100. To visualize cyst walls, rhodamine-conjugated *Dolichos biflorus* lectin (Vector Labs) was applied at 1:250 dilution overnight at 4^o^C. Cyst quantification was performed as previously described [62].

### Cell Collection and Flow Cytometry

Following euthanasia, peritoneal exudate cells (PECs) were collected from the peritoneal cavity in sterile PBS. Cells were counted using an automated cell counter, and total cell numbers were determined using the volume recovered from the peritoneum. Aliquots were used for cytospins, and stained using HEMA3 stains. For flow cytometry, cells were incubated with 1:10 FC Block (BD, 553142) for 5 minutes on ice, and subsequently incubated with fluorophore-conjugated antibodies to CD45, CD11b, CD11c, Ly6C, Ly6G. Cells were washed, resuspended in FACS buffer and samples acquired using a BD FACS Canto II flow cytometer. Analysis was conducted using Flowjo software.

### Parasite Burden and Histology

Peripheral tissues were placed in tissue lysis buffer for DNA isolation. DNA was isolated using a genomic DNA purification kit (Roche, 11796828001). To quantify parasite burden, quantitative PCR (Bioline) was conducted on isolated DNA by amplification of the *Toxoplasma* B1 gene (F: 5’ TCCCCTCTGCTGGCGAAAAGT 3’ R: 5’ AGCGTTCGTGGTCAACTATCG 3’). Parasite burden was quantified using a standard curve as previously described [63].

### Ethics Statement

All animal research was conducted in accordance with the animal welfare act, and all protocols were approved by the Institutional Animal Care and Use Committees at the University of California, Riverside (approved protocol #A-20140007) or the Indiana University School of Medicine (approved protocol #10852).

## Acknowledgements

*T. gondii* genomic and/or cDNA sequence data were accessed via http://ToxoDB.org. The authors would like to thank Drs. Elena Suvorova, Olivier Lucas, Kami Kim, Louis Weiss, and Manuel Llinas for their insights and contributions that helped this paper come together. We especially thank Dr. Erandi K. De Silva for her expertise in generating protein binding microarray results. All gene *Toxoplasma* GeneChip expression data was generated by the Moffitt Cancer Center Molecular Genomics core facility.

